# Mathematical modelling of serine integrase - mediated gene assembly

**DOI:** 10.1101/433441

**Authors:** Alexandra Pokhilko, Steven Kane, W. Marshall Stark, Sean D. Colloms

## Abstract

Site-specific recombination promoted by serine integrases can be used for ordered assembly of DNA fragments into larger arrays. When a plasmid vector is included in the assembly, the circular product DNA molecules can transform *E. coli* cells. A convenient “one-pot” method using a single integrase involves recombination between pairs of matched orthogonal attachment sites, allowing assembly of up to six DNA fragments. However, the efficiency of assembly decreases as the number of fragments increases, due to accumulation of incorrect products in which recombination has occurred between mismatched sites. Here we use mathematical modelling to analyse published experimental data for the assembly reactions and suggest potential ways to improve assembly efficiency. We assume that unproductive synaptic complexes between pairs of mismatched sites become predominant as the number and diversity of sites increase. Our modelling predicts that the proportion of correct products can be improved by raising fragment DNA concentrations and lowering plasmid vector concentration. The assembly kinetics is affected by the inactivation of integrase *in vitro*. The model also predicts that the precision might be improved by redesigning the location of attachment sites on fragments to reduce the formation of the wrong circular products. Our preliminary experimental explorations of assembly with ϕC31 integrase confirmed that assembly efficiency might be improved. However, optimization of efficiency would require more experimental work on the mechanisms of wrong product formation. The use of a more efficient integrase (such as Bxb1) might be a more promising approach to assembly optimization. The model might be easily extended for different integrases or/and different assembly strategies, such as those using multiple integrases or multiple substrate structures.

## 1. Introduction

Ordered assembly of relatively short DNA fragments into longer arrays is a critical process for many synthetic biology projects, such as installation of novel genetic circuits or metabolic pathways in host cells (1–4). Various well-established methods for DNA fragment assembly are widely used (5, 6), and recently, efficient new methods using site-specific recombination by serine integrases have been reported (7–10). Serine integrases are bacteriophage-encoded enzymes that promote recombination between sites in the phage and bacterial host chromosomal DNA. Recombination between the phage site *attP* (*P*) and the bacterial site *attB* (*B*) splices the phage DNA into the host chromosome and results in two product sites flanking the integrated phage DNA, *attL* (*L*) and *attR* (*R*), each consisting of a half-*P* and a half-*B* site (4, 11). This integrase-promoted reaction is unidirectional. However, in the presence of an additional phage-encoded protein, the recombination directionality factor (RDF), integrase promotes the reverse reaction, excision of the phage DNA from the host genome by recombination between *L* and *R* sites, thereby reconstituting *P* and *B* sites. The efficient, unidirectional nature of serine integrase-mediated recombination, along with its compact recombination sites and lack of cofactor requirements, make it attractive for many practical applications, including *in vitro* DNA fragment assembly (4, 10, 11).

In order to assemble many fragments correctly in a desired spatial order, pairs of *att* sites that are to recombine must be distinct from each other. This might be achieved by simultaneous use of several orthogonal serine integrases, but a set of such integrases is not currently available. In a published method, assembly was done using a single integrase, and pairs of *P* and *B* sites with uniquely compatible DNA sequences (10). The mechanism of serine integrase-catalysed recombination involves cleavage of the top and bottom DNA strands at the centres of the two recombining sites, followed by exchange of the cleaved ends (Fig. 1A). The top and bottom strand cleavages are separated by exactly 2 bp, so that the central 2 bp of the recombinant sites consist of a top strand (2 nt) from one of the substrate sites and a bottom strand from the other site. For successful recombination, the top and bottom strands must make correct Watson-Crick basepairs (i.e. are “matched”, Fig. 1A, B), but their sequence is not important. Compatible pairs of sites therefore have the same central 2 bp sequence. Although there are 16 possible 2-bp DNA sequences, the functional 2-fold symmetry of the *att* sites means that only 6 of these sequences can be used to specify recombination between sites in a single relative orientation (10). This method is therefore limited to assembly of up to 6 fragments in a “1-pot” reaction, although the array can be extended to larger numbers of fragments by subsequent reactions. Fig. 1C illustrates the principle of the assembly, which is based on integrase-mediated recombination between matched pairs of orthogonal *att* sites, located on linear fragments and on the plasmid vector (10).

**Figure 1.**
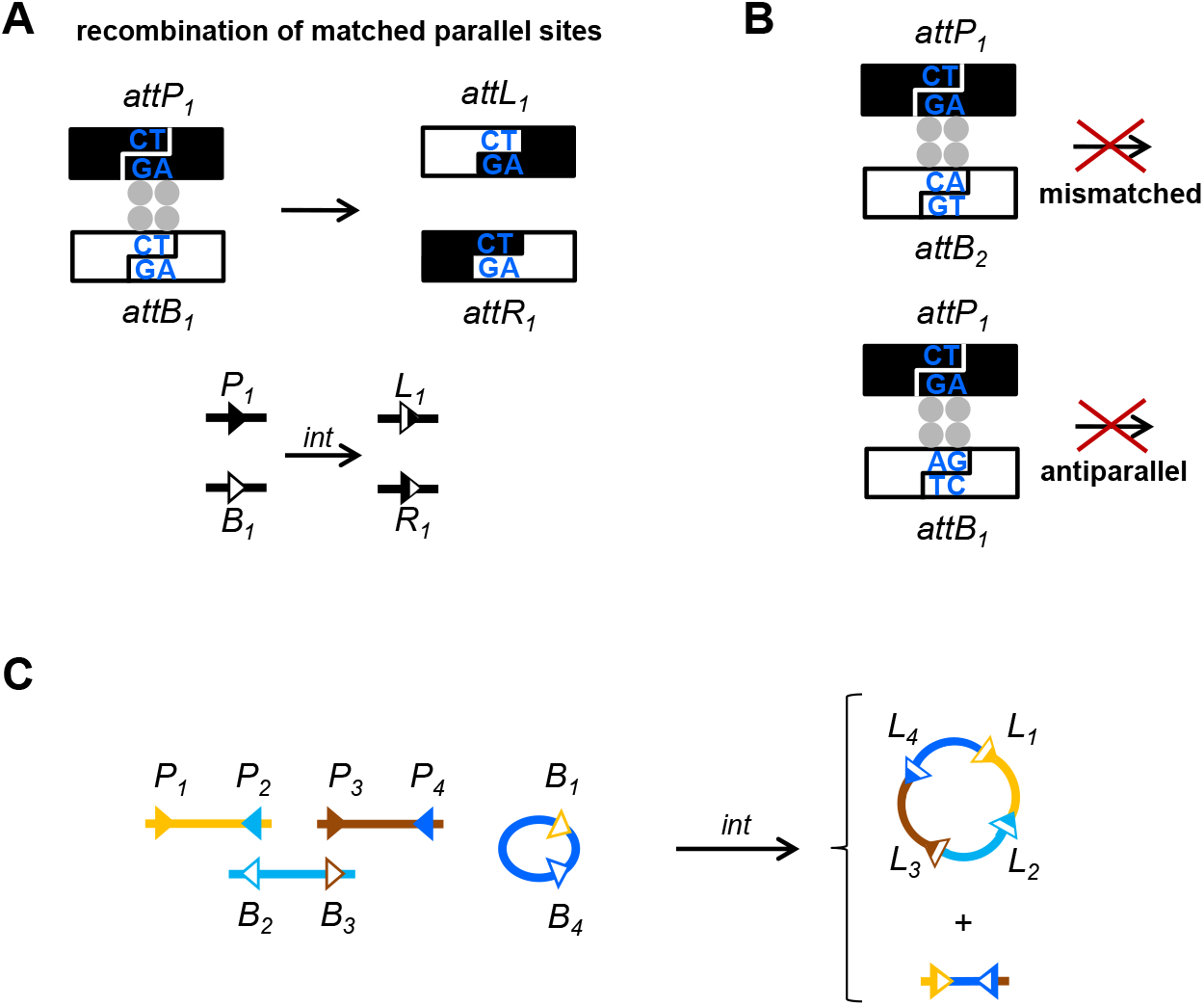
Principles of DNA assembly with serine integrase. **A**. Schematic illustration of integrase-mediated recombination between matched *attP* and *attB* attachment sites (indicated by the same index) in the parallel orientation. 4 integrase molecules (grey circles) bind DNA and form a synapse. Integrase cuts the DNA double helices at the central 2 bp of each *att* site (blue; in this example CT/AG) as shown, then rotates and re-ligates the DNA ends, resulting in the product sites *attL* and *attR*. The bottom scheme shows a compact representation of the same reaction (to be used in subsequent Figures), with sites shown by triangles (filled for *P*, empty for *B* site and half-filled for *L* and *R* sites). **B**. Examples of unproductive synapses between mismatched sites (with different central 2 bp) (top), or sites with the same central 2 bp but aligned in antiparallel (bottom). In both cases strand exchange would lead to products with mismatched basepairs, and the reaction is highly inefficient. **C**. Scheme of serine integrase-mediated gene assembly from (10) (example with 3 linear fragments). Each substrate (fragment or vector) has two sites. Matched sites are the same colour. The circular product has *L* sites between each DNA fragment. A segment of the circular plasmid substrate is excised as a linear product with *R* sites at each end. Other short fragments with *R* sites, which are created from the fragment ends by recombination, are not shown.

The method outlined above was used to assemble plasmids which could then transform *E. coli* cells and thereby introduce genes encoding metabolic pathways (10). Assembly of a plasmid encoding genes for carotenoid pigment biosynthesis gave coloured transformant *E. coli* colonies indicating production of the predicted pigment, but also many white colonies, indicating incorrect assembly (*i.e*., one or more genes missing from the plasmid). In further experiments it was observed that the fraction of incorrect products increased rapidly as the number of fragments in the construct increased, up to 82% for the assembly of 5 linear fragments (plus the plasmid vector) (10). The total yield of correct product and the rate of assembly also decreased as the number of fragments increased (observed as decreasing numbers of transformant colonies). Therefore, this assembly method requires further optimization in order to be widely applied in biotechnology. Here we use mathematical modelling of the assembly process to understand the factors affecting assembly efficiency and to explore potential ways to optimize assembly yield and accuracy. Based on the model predictions we also performed preliminary experimental tests aimed at testing predictions from the model to further explore potential approaches to assembly optimization.

## 2. Results and discussion

### 2.1. Model construction

We built a mathematical model based on the method described by Colloms *et al*. (10), in which ϕC31 integrase promoted assembly of linear DNA fragments (encoding genes for enzymes of the carotenoid biosynthetic pathway) and a supercoiled plasmid vector (see above). Supercoiled vector was used because of high recombination efficiencies with supercoiled substrates (12). All substrates (fragments and vector) have two *att* sites (as in Fig. 1C). We modelled assemblies with different type of substrates: (a) all substrates have two sites of the same type; two *P* sites (*PP*) or two *B* sites (*BB*). (b) all substrates have one *P* site and one *B* site (*PB*). (c) some substrate molecules have both sites of the same type (*PP* or *BB*) and some have one site of each type (*PB*).

Experimental monitoring of assembly efficiency was done by transforming *E. coli* cells with the assembly products and counting red, orange or yellow colonies (the colour of the pigment produced by the correctly assembled carotenoid biosynthetic pathway). The supercoiled plasmid vector has a *ccdB* gene between the two *att* sites, which is lethal in the transformed *E. coli* strain, but is deleted in correctly assembled product plasmids (10). Some incorrectly assembled cyclic products (“wrong products”, *wp*) have lost *ccdB* and can transform the *E. coli* strain, but these give rise to colourless colonies (see below).

The model incorporates several types of elementary transactions between two *att* sites. We considered only transactions between *P* and *B* sites (*P* × *B*); other possibilities (*B* × *B, P* × *P*, and all interactions involving the *L* and *R* product sites) were not included because current evidence suggests that they are weaker (13). Every reaction is assumed to start with binding of integrase to single *att* sites and the formation of a synaptic complex in which a tetramer of 4 integrase molecules connects a *P* site and a *B* site. For matched *P* and *B* sites which have synapsed in the correct (parallel) alignment, integrase quickly promotes recombination to give *L* and *R* product sites (Fig. 1A). If mismatched *P* and *B* sites form a synaptic complex or if matched sites synapse in antiparallel alignment, no products are formed (as recombination would produce mismatched basepairs in the products; see Introduction (Fig. 1B)). Accumulation of these “wrong synapses” (*ws*) is the main factor decreasing product yields in our model (14, 15), which is explained by a depletion of the substrate pools. In addition, integrase can promote recombination in these “mismatched” synaptic complexes, albeit at a relatively low rate, leading to the “wrong products” (*wp*) (16). We assumed that only a small fraction of *ws* can recombine and therefore ignored the potential depletion of the substrate pools due to *wp* formation. We consider only circular *wp* with the vector backbone which contains an origin of replication, because these *wp* can transform *E. coli*, decreasing the assembly precision. Even though the amounts of these *wp* are small (a few %), they become comparable to the amounts of correct products during assembly with large number of fragments, as discussed below.

Each elementary *P* × *B* reaction is described as a single step, leading to correct products, *ws* complexes or *wp*. Examples of the elementary reactions are shown in Fig. 2. The elementary reactions were further classified according to three characteristics (Fig. 1S): 1) the number of substrates – two (for intermolecular reactions) or one (for intramolecular reactions); 2) type of substrate(s) (linear or circular supercoiled); 3) relationship of the interacting sites – matched or mismatched (including antiparallel). All recombination reactions are assumed to be irreversible, whereas *ws* formation is assumed to be reversible (Fig. 2, Fig. 1S). The model also describes the inactivation of integrase observed *in vitro* (possibly due to precipitation) (12) and protection of integrase against inactivation upon its binding to DNA (section 3).

Assembly involves sequential recombination reactions which join substrates together (Fig. 3). The number of steps increases with the number of substrates. In the following discussion, we use the term “fragment” to mean a linear DNA molecule being assembled with *att* sites at each end. Assembly starts with recombination between two sites located on free substrates; one a circular supercoiled plasmid and the other a linear fragment. We assume that the plasmid supercoiling is released after the first recombination event (16), giving a linear intermediate as illustrated in Fig. 3A. In subsequent steps, the intermediates are extended by recombination with further fragments. The concentrations of free (unbound) fragments are larger than the concentrations of the longer recombined intermediates in our model. Therefore, we assumed for simplicity that the intermediates are only elongated by recombination with free fragments. A final, intramolecular recombination event creates the assembled circular product ***L_fin_**. CcdB* is excised from the vector during the assembly (Fig. 3A, B).

In addition to the correct product ***L_fin_***, circular *wp* are formed (Fig. 3B). Three types of *wp* were considered in our model based on previous observations (10). The first type, *wp_0_*, results from intramolecular recombination of the mismatched sites of a *PB* vector, with no fragments involved (Fig. 3B, (10)). The second type, *wp_1_* results from recombination between the vector and a single fragment with one matched and one mismatched site (Fig. 3C, Fig. 2S C). This is the main type of *wp* observed during *PP/BB*-type assembly (10), as we discuss below. A third type of *wp* (*wp_2_*) results from intramolecular recombination of mismatched sites in longer intermediates consisting of two or more fragments (Fig. 2S D). We assumed for simplicity that formation of *wp_1_* and *wp_2_* starts from correct recombination between matched sites, followed by wrong intramolecular recombination of the mismatched sites (Fig. 3C, Fig. 2S).

The model parameters (Table 1 of the Appendix A) were fitted to the published data on accumulation of correct (*L_fin_*) and wrong (*wp*) products (10). The model explains the key observations and suggests potential ways of optimizing assembly, as discussed below.

**Table 1.** The values of the model parameters.

|  |  |  |  |  |  |
| --- | --- | --- | --- | --- | --- |
| parameter | $K_{bl}$ | $k_{+b}$ | $k_{-b}$ | $k_{ia}$ | $k$ |
| value | 0.1 $\mu\text{M}$ | 60 $\text{min}^{-1}\mu\text{M}^{-1}$ | 0.6 $\text{min}^{-1}$ | 0.01 $\text{min}^{-1}$ | 8 |
| ref. | (12) | (12) | fit | fit | fit |
| parameter | $k_{+r}$ | $k_{+r0}$ | $k_{+s}$ | $k_{+s0}$ | $k_{-s}$ |
| value | 0.3 $\text{min}^{-1}$ | 2.3 $\text{min}^{-1}\mu\text{M}^{-1}$ | 0.001 $\text{min}^{-1}$ | 0.1 $\text{min}^{-1}\mu\text{M}^{-1}$ | 0.006 $\text{h}^{-1}$ |
| ref. | fit | fit | fit | fit | fit |
| parameter | $k_{+rw0}$ | $k_{+rl}$ | $k_{+rw}$ | $k_{ia}$ | $k$ |
| value | 0.001 $\text{min}^{-1}$ | 0.32 $\text{min}^{-1}\mu\text{M}^{-1}$ | 0.00008 $\text{min}^{-1}$ | | |
| ref. | fit | fit | fit | fit | fit |

**Figure 2.**
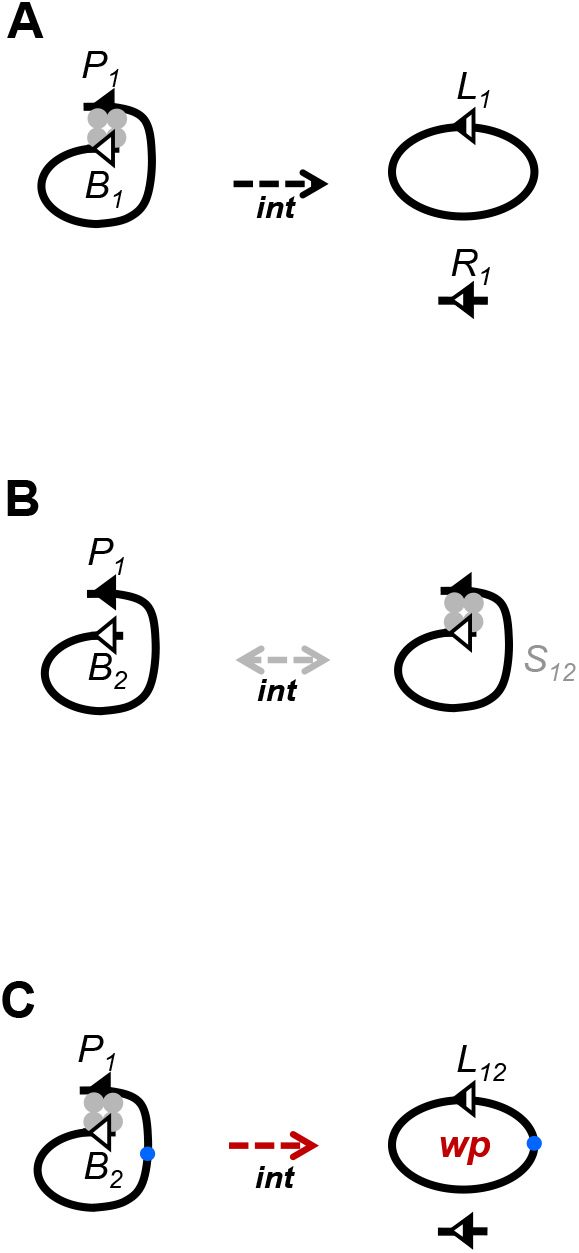
Three types of elementary reactions between *P* and *B* sites included in the model. Matched sites are indicated by the same indices (*e.g., P_1_, B_1_*). **A**. An example of correct recombination. Synapsis between matched *P* and *B* sites mediated by integrase (grey circles) leads to recombination product sites *L* and *R*. **B**. An example of the formation of wrong synapse *ws* (named S_12_) between mismatched sites *P_1_* and *B_2_*. **C**. An example of wrong recombination between mismatched sites, leading to circular *wp*. Only *wp* bearing the origin of replication (blue dot) were considered. Recombination reactions are assumed to be irreversible and formation of *ws* is reversible.

**Figure 3.**
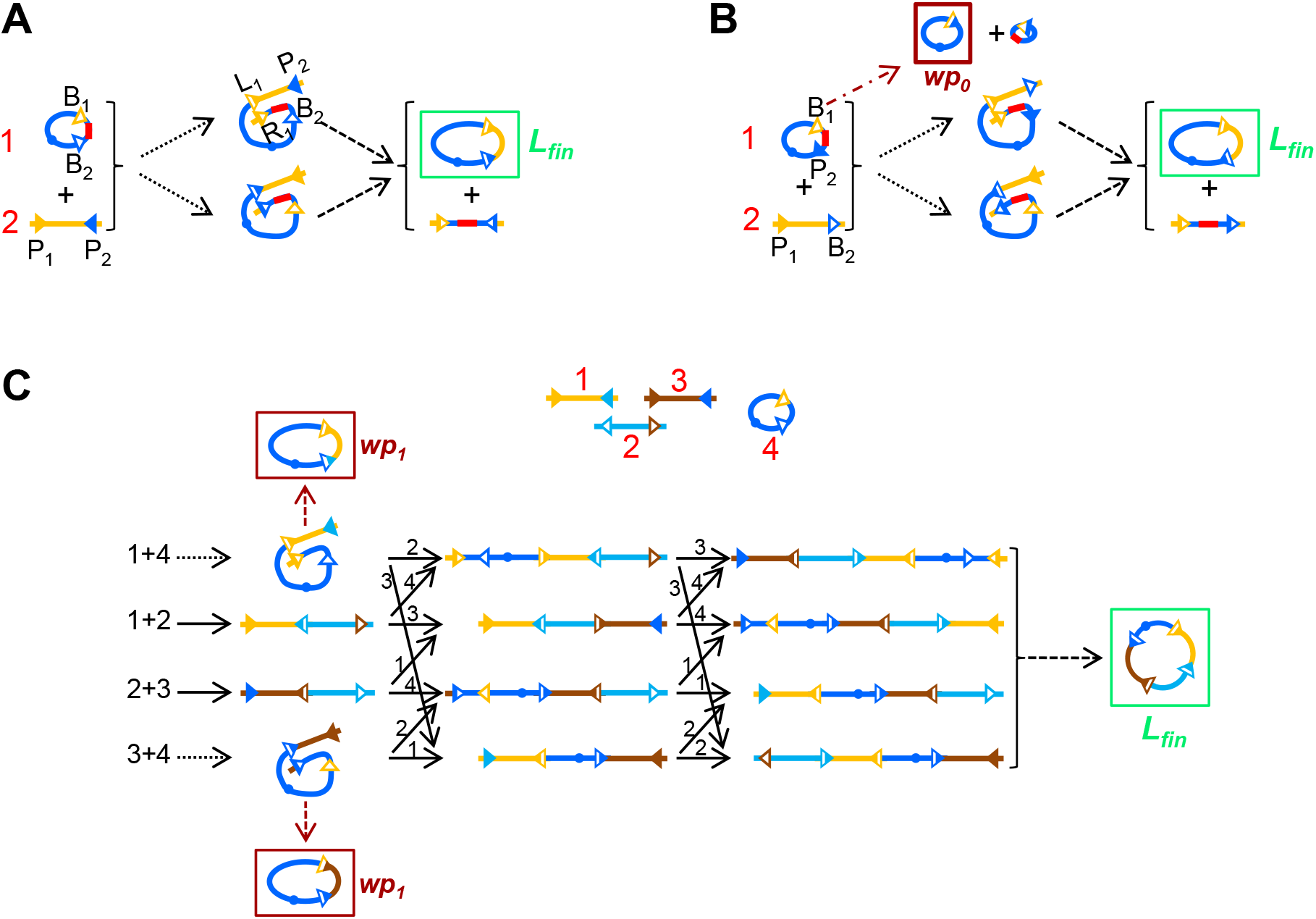
Model schemes for gene assembly as tested experimentally in (10). The substrates (fragments and vector) are marked by red numbers. The orientation of the sites on substrates corresponds to (10) (Fig. 1A). **A**. A 1-fragment assembly with *PP/BB*-type substrates. **B**. A 1-fragment assembly (insertion of a single fragment into the vector) with *PB* substrates. **C**. 3-fragment assembly with *PP/BB*-type substrates. The model describes several ways for assembly, depending on which substrates recombine first. The *ccdB* gene is shown by a red line (on A, B only). Short recombination products with *R* sites are not shown (except for the *ccdB* gene in A, B). Black numbers before/above the reaction arrows refer to the interacting species. Different types of arrows show different types of elementary reactions (as in Fig. 1S): The formation of circular wrong products *wp* bearing the origin of replication (blue dots) is shown by red arrows. *Wp* are classified into *w_0_, wp_1_* depending on the number of fragments involved: *w_0_* consists only of the *PB* vector (B); *wp_1_* include a single fragment with one matched and one mismatched site to the vector sites (C, D). The complete schemes, including formation of *ws* and 5- fragment assembly, are shown in Fig. S1.

### 2.2. Kinetics of assembly. Synapsis between mismatched sites decreases yields and increases mistakes in multi-fragment assemblies

Two important characteristics of the assembly reactions are: the **yield** (total amount of products relative to the theoretical maximum) and the **precision** (% of the total number of circular products that are correctly assembled). High yields might be especially important for combinatorial assemblies with multiple versions of substrates, while high precision might be particularly important for assemblies with unique fragments, such as the ones described here. In our model, yield corresponds to the measured total number of transformed colonies (normalized to an experimentally determined maximum value), and the precision corresponds to % of correct colonies (10).

In our model, the product yields and the assembly rates decrease with increasing number of fragments, in agreement with the data (Fig. 4A-C, (10)). Thus, the simulated kinetics of 1-fragment assembly (insertion of a single fragment into the vector) is fast (half-saturation time of the product formation, T_0.5_ < 1 h) and the product yield is high (80%) (Fig. 4A). The increased accumulation of *wp* during *PB*-type assembly relative to *PP/BB* assembly (Fig. 4A, (10)) is explained by intramolecular recombination of mismatched vector sites yielding *wp_0_* (Fig. 3B). For 3-fragment assembly the kinetics are slower (T_0.5_ ∼5 h) and product yields are lower (10%) than for 1-fragment assembly (Fig. 4B). These effects become even more pronounced in simulations of 5-fragment assembly, with T_0.5_ ∼ 12 h and 4% yield (Fig. 4C). The model explains the reduced product yield by increased formation of non-productive synaptic complexes *ws* (14, 15), which depletes the concentrations of free fragments (Fig. 4D, E) and slows down assembly. In our model the precision (% of correct product after 24 h of reaction) decreases with increase of the number of fragments (10), from 100% for 1-fragment *PP/BB* assembly (Fig. 4A) to 83% and 17% for 3- and 5-fragment assemblies, in agreement with the data (Fig. 4B, C, (10)). This decrease of precision is due to increased amounts of circular *wp* relative to *L_fin_* (Fig. 4A-C). In our simulations of *PP/BB* assembly, most *wp* are represented by *wp_1_* (Fig. 4C, Fig. 3C, Fig. 2S, (10)), consisting of the vector and a single fragment. The concentrations of *wp_2_* (with more than one fragment) are low in our model due to reduced concentration of longer intermediates. The mechanism of *wp_1_* formation in our model involves reaction between the vector and a fragment with one site matching the vector and one mismatched site (Fig. 2S C, D). *Wp_1_* are formed in this way during 3- and 5-fragment assemblies, but the absolute levels of *wp_1_* are higher in 5-fragment assembly than in 3-fragment assembly, in agreement with the data (Fig. 4B, C, (10)). This is explained in our model by slower inactivation of integrase during 5-fragment assembly (Fig. 4F) due to its protection by larger amounts of DNA. Slower inactivation of integrase leads to prolonged accumulation of both correct and wrong products, resulting in higher absolute values of *wp* (Fig. 4C).

**Figure 4.**
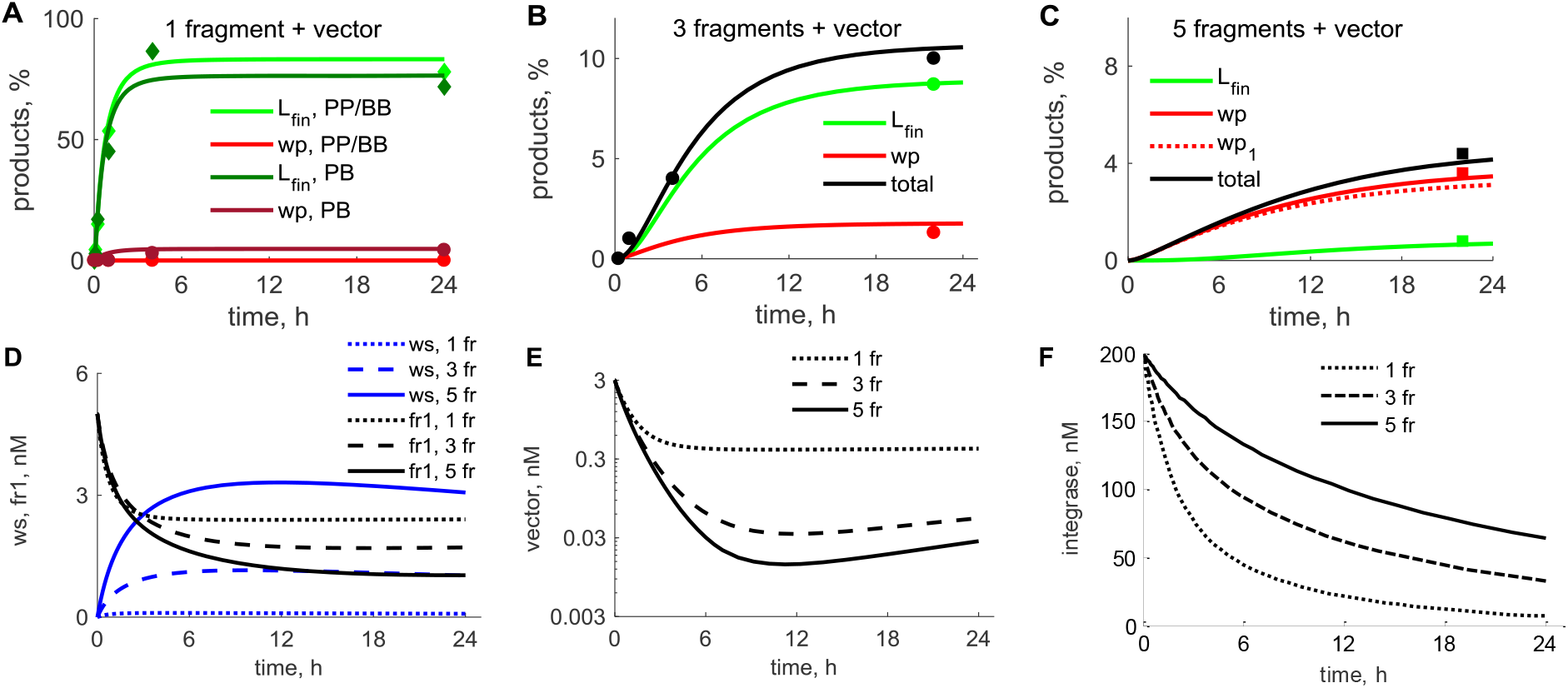
Simulations of experimental kinetics of 1-, 3- and 5-fragment assembly. Modelling is shown by lines and data (10) by symbols. **A**. Kinetics of the correct (*L_fin_*, green) and wrong (*wp_0_*, red) products during 1-fragment assembly. Lighter green/red lines and symbols correspond to *PP/BB*-type assembly, and darker lines/symbols correspond to *PB*-type assembly. **B, C**. Kinetics of *L_fin_* (green), *wp* (red solid line), and total products (*L_fin_* + *wp*, black) during 3-fragment assembly (B) and 5-fragment assembly (C) with *PP/BB* fragments. The major *wp_1_* fraction of *wp* is shown by the red dotted line in C; the remaining *wp* is *wp_2_*. **D** shows the concentrations of the wrong synapses (*ws*, blue) and fragment 1 (black). **E** shows the concentrations of the vector. **F** shows the amounts of active integrase. Dotted, dashed and solid lines on D, E, F correspond to 1-, 3- and 5-fragment *PP/BB* assemblies, respectively. Initial concentrations of fragments were 5 nM each, vector (plasmid) was 3 nM and integrase was 200 nM (10).

### 2.3 Optimization of substrate concentrations increases L_*fin*_/wp ratio

Next, we used the model to explore the dependence of *L_fin_* and *wp* levels on substrate concentrations. Fig. 5A-C show that decrease of vector concentration and increase of fragment concentrations is beneficial for assembly because these changes increase the *L_fin_*/*wp* ratio. For example, 2-fold increase of fragment and 3-fold decrease of the vector concentrations relative to the control experiments (10) is predicted to cause 3-fold increase of the % of *L_fin_* from maximal amount of the cyclic products and 1.2-fold increase of the % of *wp*, leading to 2.5-fold increase of *L_fin_*/*wp* (Fig. 5D). Increased fragment concentrations increase the % of *L_fin_* due to acceleration of initial recombination steps. Higher DNA levels (i.e. higher fragment concentrations) slow down inactivation of integrase, resulting in prolonged accumulation of both *L_fin_* and *wp* (Fig. 5D). We conclude that optimization of substrate concentrations might increase *L_fin_*/*wp* ratio, but does not reduce the *wp* content relative to maximal amount of the cyclic products.

**Figure 5.**
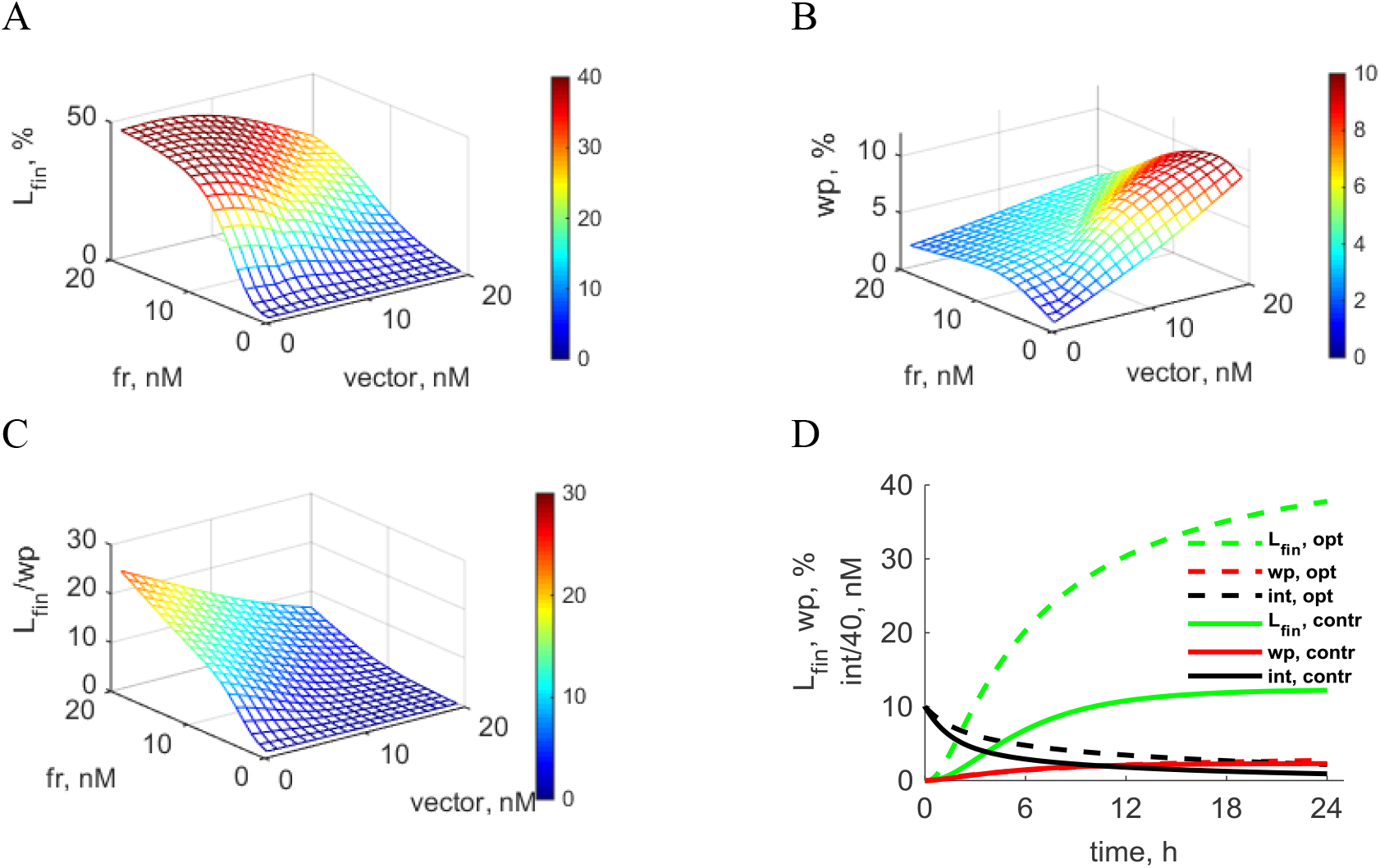
Dependence of the efficiency of 3-fragment *PP/BB*-type assembly on the concentrations of DNA substrates. **A**. Correct product *L_fin_* (% of maximal amount of cyclic products). **B**. % of the wrong products *wp*. **C**. *L_fin_*/*wp* ratio. Calculations are for 24-h reactions. **D**. The kinetics of the % of *L_fin_* (green), % of *wp* (red) and concentration of integrase (int, black) for the control (solid lines) and optimized (dashed lines) conditions. The % of *wp* is nearly identical in control and optimized conditions. Concentrations are: control, 5 nM each fragment, 3 nM vector (plasmid); optimized conditions,10 nM each fragment, 1 nM vector. Initial concentration of integrase is 400 nM in all simulations. Integrase concentrations are scaled to fit to the figure.

### 2.4. Wrong product reduction using mixed assembly of PB and PP fragments, and BB vector

Model simulations demonstrate that the efficiency of assembly is lower when all the fragments are *PB* than when mixtures of *PP* and *BB* fragments are used (Fig. S3A). Even for 1-fragment *PB* assembly, the precision is lower (94% after 24 h) compared to 100% for *PP/BB* assembly (Fig. 4A). The precision further decreases for 3-fragment assembly, from 83% for *PP/BB* assembly (Fig. 4B) to 46 % for *PB* assembly (Fig. S3B). This decrease corresponds to increased *wp* and decreased *L_fin_* (Fig. S3 B). The increase in *wp* during *PB* assembly is explained by high intramolecular recombination of the mismatched sites of the *PB* vector, leading to the accumulation of *wp_0_* (Fig. S3 A, B, (10)). We conclude that the use of a *PB* vector in gene assembly should be avoided.

For *PP/BB* assembly, *wp* are mainly represented by *wp_1_* (see above). However, an assembly can be designed by using a *BB* vector and *PB* fragments which cannot form *wp_1_* (Fig. 6A). The single *PP* fragment which is required for this assembly (and which could potentially form *wp_1_*) might be placed in the middle of the array (fragment 2 in Fig. 6A), so that it has two mismatched sites with the vector. In this assembly design only *wp_2_* (with more than one fragment) are possible (Fig. 6A). Modelling of this assembly type predicts a 4-fold decrease of the absolute amounts of *wp* compared to *PP/BB* assembly, resulting in increase of the precision from 84% to 96% of the correct product after 24 h (Fig. 6B). The model further predicts that a 7-fold decrease of *wp* (Fig. 6C) can be achieved by preincubation of the reaction without fragment 2. The decrease of *wp* is explained by consumption of the vector by productive recombination reactions during preincubation (Fig. 6C).

**Figure 6.**
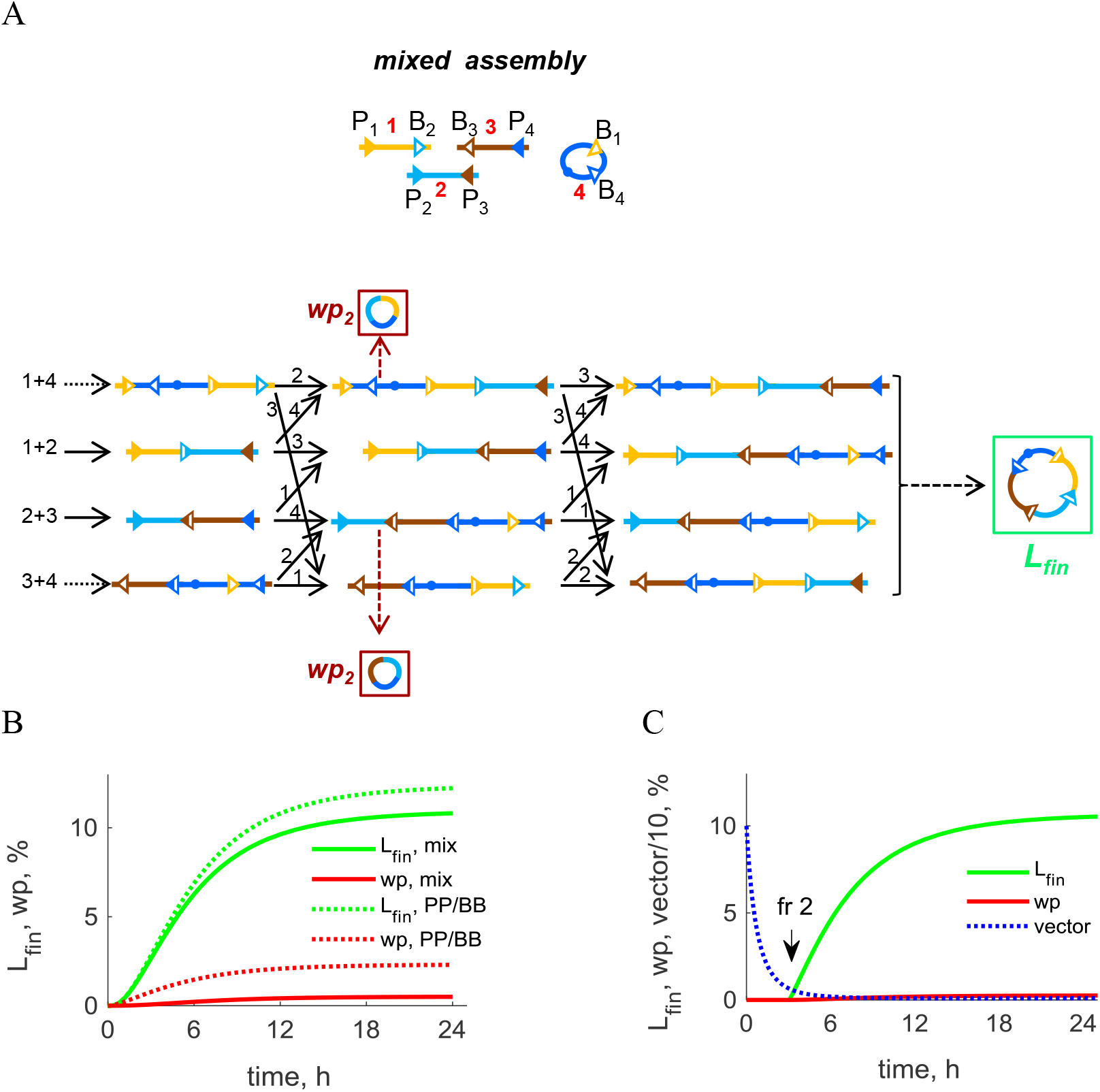
Simulations of 3-fragment assembly with mixed types of fragments. **A**. Model scheme. The assembly includes a *BB* vector, two *PB* fragments and one *PP* fragment (fragment 2). The complete scheme (including formation of *ws*) is shown in Fig. S4. **B**. Comparison of *L_fin_* (green) and *wp* (red) kinetics during *PP/BB* assembly (dotted lines) and mixed assembly (solid lines). **C**. Kinetics of *L_fin_* (green), *wp* (red) and vector (blue) after 3 h preincubation of all substrates (excluding fragment 2) with integrase. The time of the addition of fragment 2 is shown by the arrow. B, C. Concentrations of fragments were 5 nM, vector 3 nM and integrase 400 nM. The levels of all variables are shown as % of maximal amount of product (3 nM).

A similar approach can be applied for optimization of 5-fragment assembly, using fragments of the *PB* type except for a single *PP* fragment in the middle of the assembly (Fig. S5). The model predicts a 17-fold decrease in the absolute amount of *wp* compared to *PP/BB* assembly (Fig. 7B), resulting in increase of the precision from 19% for *PP/BB* assembly to 72% for mixed fragment assembly (Fig. 7). The improvement is explained by the absence of *wp_1_* products.

In summary, assembly using *PB* fragments except for a central *PP* fragment may be a good strategy to improve assembly precision. However, the relatively low precision of ϕC31 integrase (large number of *wp*) suggests that using a more efficient integrase (such as Bxb1, (16)) and/or multiple orthogonal integrases might be a better strategy, as further discussed below.

**Figure 7.**
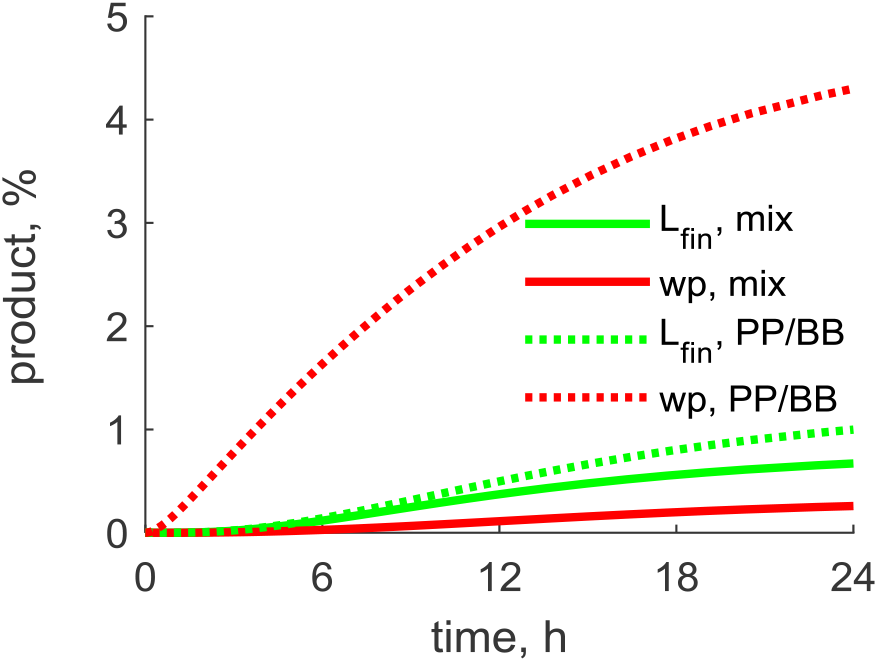
Comparison of the kinetics of 5-fragment assembly with mixed fragment type and *PP/BB* assembly. The mixed-fragment assembly uses fragments of the *PB* type except for one *PP* fragment in the middle of the array and the *BB* vector (Fig. S6). The kinetics of *L_fin_*(green) and *wp* (red) fractions is shown for the mixed-fragment assembly (solid lines) and for *PP/BB* assembly (dotted lines). Simulations used concentrations as in Figure 6.

### 2.5 Preliminary experimental results

Motivated by our model predictions, we did preliminary experimental tests of the assembly reactions. We first performed 1-fragment assembly using *BB* vector and a *PP* fragment with one matched and one mismatched site to the vector sites (Fig. 8A). This assembly does not result in the correct cyclic product, but only *wp*. We used the same conditions as in (10), with TT and TC central dinucleotides in the vector sites and TT dinucleotide in the matched fragment site. For the mismatched site we considered 4 versions of the central dinucleotides: CA, CC, CT, GT. The fragment with two matched sites (TT and TC) was used as a control, to compare the production of correct and wrong products. Fig. 8B shows the kinetics of *wp* formation for 4 mismatched fragments and *L_fin_* kinetics for the matched fragment. These data suggest that, although the *wp* kinetics might vary depending on the sequences of the central dinucleotides, the *L_fin_* / *wp* ratio is around 20–100. This is ∼10-fold lower that in the model (Fig. 8C). A potential reason for the observed high levels of *wp* might be the production of *wp* by the transformed cells from the linear product of recombination between matched sites of the fragment and vector (product 3 on Fig. 8A). The level of this intermediate is expected to be high, because it cannot be used for production of the correct product. This possibility might be investigated by transforming cells with the respective linear products.

**Figure 8.**
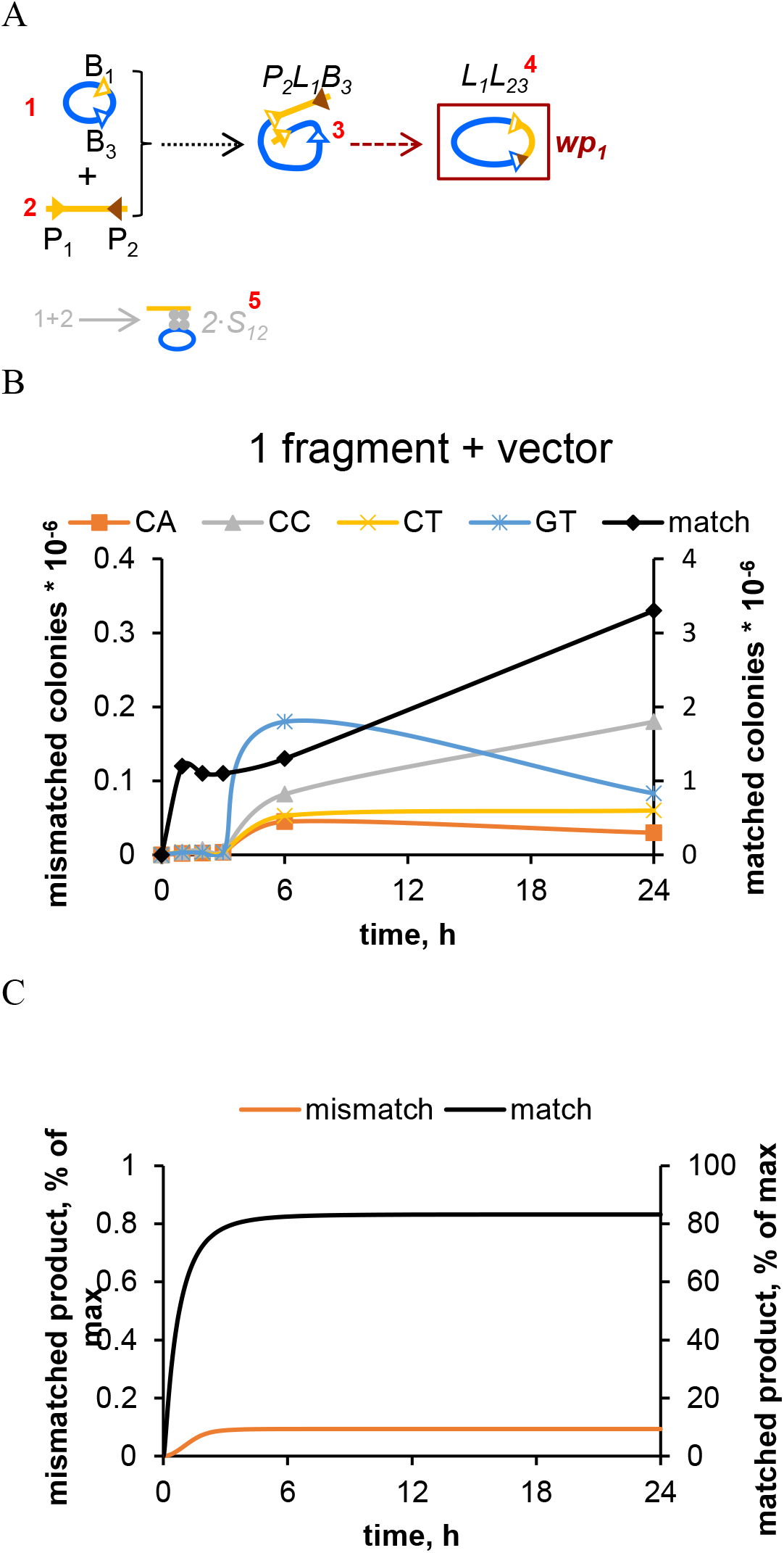
The kinetics of *wp* formation in 1-fragment assembly between the vector and a fragment with one mismatched site. A. Scheme of the assembly reactions with a fragment having one matched and one mismatched site to the vector sites. B. Experimental assembly kinetics, measured as the number of colonies with circular products. The vector has TT and TC central dinucleotides, while the fragment has a TT dinucleotide in the matched site and various dinucleotides (CA, CC, CT, GT, as indicated in the legend) in the mismatched site. The control kinetics of the correct *L_fin_* product, resulting from assembly with a fragment with matched sites (TT, TC dinucleotides) is shown by the black line. Initial concentrations of fragments were 5 nM, vector was 3 nM and integrase was 200 nM. C. The model simulations of the data shown in B. The concentrations of products are normalized to the vector concentration (3 nM).

We next experimentally compared *PP/BB, PB* and mixed 3-fragment assembly of the lycopene synthesis pathway (see ref. 10). All reactions were for 6 h, and the final amounts of the correct (red) and wrong (white) colonies were measured (Table 2). The mixed-fragment assembly was carried out with a *PP* fragment being placed at the end of the array (Fig. 9). Precision of the *PB* assembly was lower than for the *PP/BB* assembly, as expected (10) (lines 1,2 of Table 2). Unexpectedly, the mixed-fragment assembly was found to have ∼2-fold lower precision than *PP/BB* assembly (compare lines 2 and 3). Adding more integrase in the middle of the assembly reaction (at 3 h) improves the mixed-fragment assembly precision 1.7-fold (lines 3,4), supporting the idea that integrase inactivation affects assembly. However, the delayed addition of *PP* fragment (together with integrase) decreased the precision (lines 6), contrary to the model prediction. We also observed this for the 3 h (but not 20 min) addition of *PP* fragment during *PP/BB* assembly, as discussed below (lines 8,9, Table 3). We also tested heating the reaction mixture before adding more integrase, which was expected to release the substrates and integrase from *ws* and thus increase precision. However, the heating reversed the effect of integrase addition (lines 5,4). The reasons for this effect need to be investigated further. In particular, the possibility of cell transformation-related artefacts needs to be excluded.

**Table 2.** Comparison of the total product (colony count) and % correct product (% of red colonies, producing lycopene product) for different assembly types. The assemblies were started from mixtures of the indicated fragments (5 nM each), 3 nM vector (*BB*-pSira1 or *PB-* pSira3) and 200 nM of integrase. After 3 h some samples received addition of another 200 nM integrase (int), either alone (lines 4,5) or with *PP* fragment (lines 6,7). Lines 5,7; samples were heated after 3 h for 10 min at 80° C before addition of more integrase and *PP* fragment. All reactions were performed on the same day. See Methods for more details.

| reaction N | starting reaction mix | delayed additions | colony count (per ml) | % correct |
| --- | --- | --- | --- | --- |
| 1 | PB/PB/PB+pSira3 | 0 | $3.8 \cdot 10^5$ | 0 |
| 2 | PP/BB/PP+pSira1 | 0 | $1.5 \cdot 10^5$ | 66 |
| 3 | PB/PB/PP+pSira1 | 0 | $1.7 \cdot 10^5$ | 29 |
| 4 | PB/PB/PP+pSira1 | 3 h + int | $2.3 \cdot 10^5$ | 48 |
| 5 | PB/PB/PP+pSira1 | 3 h 80° C + int | $2 \cdot 10^5$ | 29 |
| 6 | PB/PB+pSira1 | 3 h + PP + int | $5 \cdot 10^4$ | 17 |
| 7 | PB/PB+pSira1 | 3 hr80° C + PP + int | $2.4 \cdot 10^4$ | 39 |

**Figure 9.**
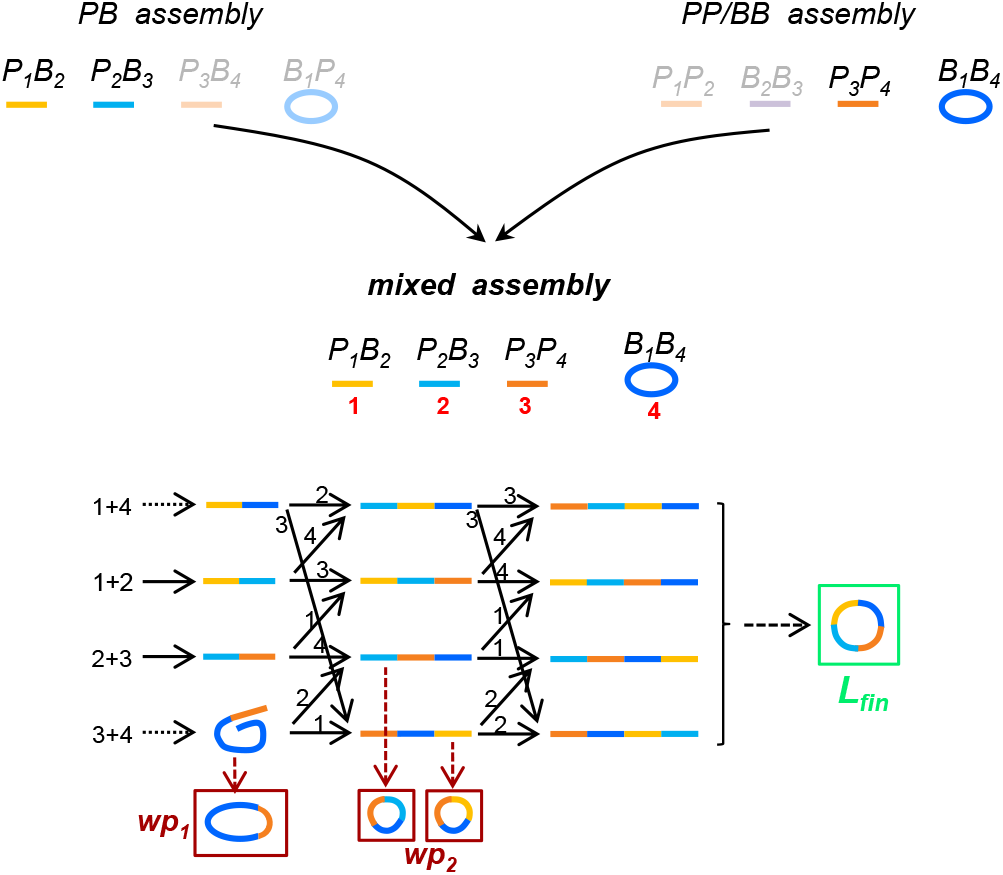
Scheme for 3-fragment assembly with mixed types of fragments. The top part of the figure shows the construction of mixed-fragment assembly from the previously developed *PB* and *PP/BB* assemblies: the first two fragments of the mixed-fragment assembly are taken from the *PB* assembly, while the last fragment and the vector are taken from the *PP/BB* assembly. The assembly includes 2 fragments of the *PB* type (from the *PB* assembly), one *PP* fragment (fragment 3) and the *BB* vector (from the *PP/BB* assembly). The designations are the same as in Fig. 3.

Another potential reason of the negative effect of delayed addition of the *PP* fragment in the mixed assembly (line 6) might be related to the specific sequence of the central dinucleotides. For all our experiments we used the same sequence as in (10): TT-CT for fragment 1; CT-GT for fragment 2; GT-TC for fragment 3; TT-TC for the vector (*BB* vector called pSira1; *PB* vector called pSira3). Therefore, the *PP* fragment has GT dinucleotides, which seems to lead to highest *wp* after 6 hours of the assembly (Fig. 8B). Moreover, the *PP* fragment was located at the end of the assembly, but not in the middle, as was suggested by the model (Fig. 9A, Fig. 6A), increasing *wp*.

After these optimizations and tests, the model parameters need to be adjusted to the new data. In particular, the model parameters were fitted to published data using approximate estimations of the maximal number of colonies (10). However, more precise calibration of the number of colonies, corresponding to the known amount of product, is necessary for parameter optimization. Our preliminary titration of the number of colonies with the known amounts of the final product gives ∼10-fold lower estimates for the total amount of product (Fig. 11). This suggests that the absolute amount of products might be over-estimated in the model, which would result in under-estimation of the *ws* concentrations.

To explore the *wp* composition, we extracted DNA from cells transformed with the products of the reaction from Table 2, and performed agarose gel electrophoresis. The products observed (Fig. 10) include *wp* having a single *PP* fragment (1 or 3, called *wp_1_* in the model), or two fragments (1+3 or 2+3, called *wp_2_* in the model) and the correct *L_fin_* product (1+2+3). The amount of *wp_2_* seems to be higher that is predicted by the model, which might explain the low efficiency of the mixed-fragment assembly.

**Figure 10.**
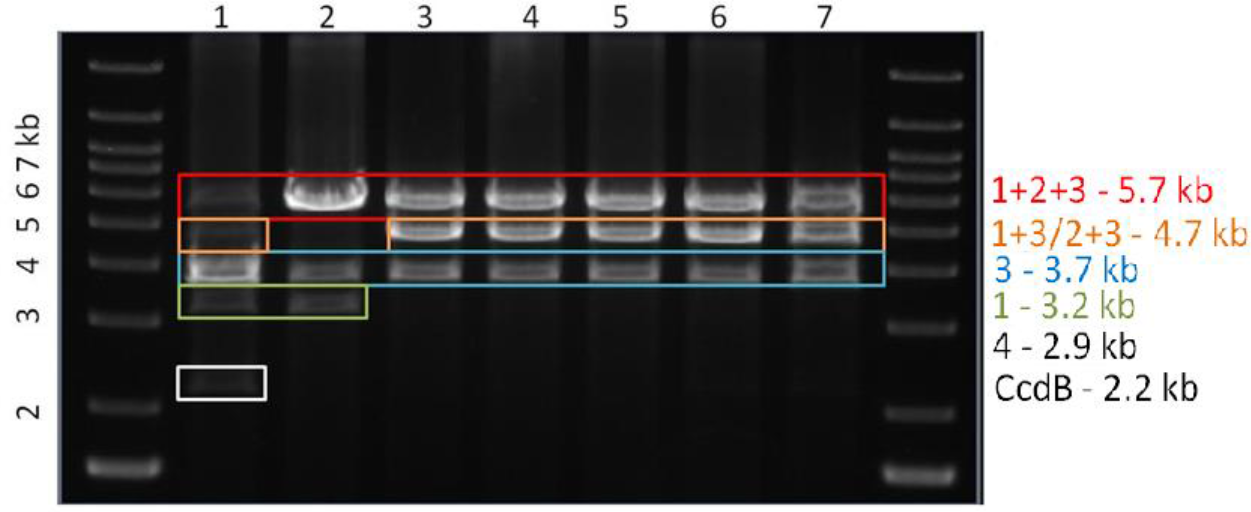
Gel electrophoresis of the restriction enzyme-linearized products of reactions from Table 2. Cells were transformed with the reaction products for 1 hour and then grown on a plate overnight. Lines 1–7 correspond to reactions of Table 2. The colours mark circular *wp* with fragment 1 (green), 3 (blue), 1+3 or 2+3 (orange); 1+2+3 (red, correct product) and the vector 4 (black). The location of the fragment containing the *ccdB* gene is also shown.

Our preliminary conclusion is that ϕC31 integrase seems to produce a lot of *wp* under various conditions. Even though assembly might be optimized by using integrase addition and heating, the choice of integrase might need to be reconsidered. Even though ϕC31 integrase is the most widely used, some other integrases, such as Bxb1 (16) might be more appropriate for gene assembly.

## 3. Methods

### 3.1. Model schemes and equations

The model construction is described in Results. The elementary reactions of *P* and *B* sites result in the correct recombinant products (Fig. S1 A, C); wrong synapses, *ws* (Fig. S1 B, D) and wrong products, *wp* (Fig. S1 E, F). The intermolecular reactions are between two linear fragments or between linear and circular substrates (Fig. S1). Cross-reactions between vector molecules (dimerization) were ignored due to their lower probability (12) and expected depletion of the vector concentrations during the assembly. The kinetics of the reactions between matched sites is modelled as independent of the exact sequence of the central 2 bp (10). Due to the absence of data, we simply assumed that the kinetics of all elementary reactions do not depend on the sequence of the central dinucleotide.

Fig. S2 shows detailed reaction schemes of the model for 1-, 3- and 5-fragment assembly of the *PP/BB* type (Fig. S2 A, C, D) and for 1-fragment assembly of the *PB* type (Fig. S2 B). The species names indicate the presence of particular sites, *e.g*., the *P_2_L_1_B_2_* species has free *P_2_, B_2_* sites able to react further, and an *L_1_* site resulting from *P_1_* + *B_1_* recombination. The first steps of the assembly give products of recombination between a fragment and the vector (*e.g*., for *P_2_L_1_B_4_*, Fig. S2 C) or between two fragments (*e.g., P_1_L_2_B_3_*, Fig. S2 C). The first recombination step between the supercoiled vector and a fragment was assumed to be faster compared to intermolecular recombination reactions between linear substrates (12). We assumed that the rates of all intermolecular reactions between linear substrates are independent of the fragment lengths, for relatively long DNA fragments used in the gene assembly (10, 17). Additionally, circular *wp* are produced by intramolecular recombination between mismatched *P* and *B* sites in the parallel orientation (Fig S1 E, F). In our model, we counted only *wp* resulting from a single wrong recombination, and ignored the *wp* which might result from two or more wrong recombination evens, due to their low probability.

In addition to recombination reactions, the model describes the formation of *ws* between mismatched *P* and *B* sites. We included only the most abundant *ws* formed between the sites of free substrates, which was sufficient to describe the data. The *ws* formed during reactions with the same substrates but different sites are described as a single entity, for simplicity. Depending on a fragment type (*PP/BB* or *PB*), the formation of *ws* might involve 1 or 2 sites of each fragment, which for the latter case is reflected by the corresponding number 2 before the *ws* species on Fig. S2. For example, the formation of *ws* named *S_12_* can involve both *B* sites of the vector (number 1 on Fig. S2 A) and both *P* sites of the fragment 2, which is reflected by a coefficient 2 before *S_12_*. However, for mixed-fragment assembly (*PP/BB* and *PB* fragments mixed together), the formation of *ws* sometimes involves only one site of a fragment. For example, only the *B* site of a *PB* fragment can form *ws* with a *PP* fragment, which would be reflected by an absence of coefficient 2 before the respective *ws* species on the schemes. *PB* assembly also includes intramolecular formation of *ws* due to reactions between mismatched *P* and *B* sites located on the same fragment (*e.g., S_P1B2_*, Fig. S2 B) or vector (*S_P2B1_*, Fig. S2 B).

The model also describes the inactivation of integrase, possibly due to its precipitation, which is observed *in vitro* (12). Inactivation of integrase is relatively quick (∼ hours), but it slows down in the presence of the DNA due to the binding of integrase to the DNA (12). The observations on the prolonged kinetics of 5-fragment assembly, with higher accumulation of *wp* (Fig. 4C) suggest that integrase might be inactivated more slowly during assemblies with more fragments, possibly due to the larger amount of the total DNA. This was described in the model through protection of free integrase from inactivation upon its binding to DNA. The total binding sites on DNA (*B_0_*) were assumed to be proportional to total DNA concentration (*D*): *B_0_ = k·D*. The total concentration of active integrase is described as a sum of free and bound integrase concentrations, *int_tot_ = int_free_ + int_bound_*. The equations describing integrase inactivation are: 
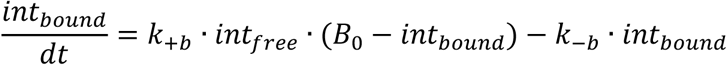

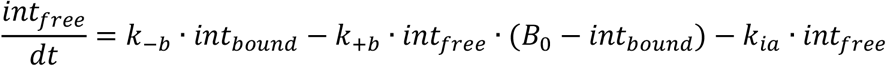

Where *int_free_* and *int_bound_* are concentrations of free and bound integrase, *B_0_* and (*B_0_ – int_bound_*) are the concentration of total and free binding sites. In section 2 we show that integrase inactivation is expected to have a noticeable effect on the assembly kinetics.

Based on tetramerization of integrase and threshold dependence of reactions on integrase concentration (12), we assumed that reaction rates depend on integrase concentration in a cooperative manner. Therefore, all reaction rates were multiplied by a function *int4(t)*: 
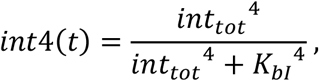
where *K_bI_* = 100 nM is integrase concentration for half-maximal rate, with the working range of integrase concentrations being typically between 200 and 400 nM (12). In most simulations we use an integrase concentration of 400 nM. In simulation of the (10) data we use 200 nM of integrase.

The model parameters were collectively fitted to the data on *L_fin_* and *wp* kinetics during 1-, 3- and 5-gene assembly of fragments encoding enzymes of the carotenoid biosynthesis pathway (Fig. 4C). The experimental levels of *L_fin_* and *wp* were assumed to be proportional to the number of correct (coloured, for 3- and 5-gene assembly, or antibiotic-resistant for 1-gene assembly) and wrong (white) colonies of transformed *E. coli* cells (normalized to maximum based on (10)). In our model the rate constants mostly affect the kinetics of the products and dissociation constants mostly affect their final levels. The parameters of *wp_0_* formation were fitted to the data on *PB* assembly, while the parameters of *wp_1_* / *wp_2_* formation were fitted to the data on *PP/BB* assembly (Fig. 4A-C). The parameters of correct recombination between a fragment and the vector were fitted to the data on higher efficiency of 1-fragment assembly.

The full set of model equations is presented in the Appendix A, together with the parameter values (Table 1). The system of ODEs was solved using MATLAB, integrated with the stiff solver ode15s (The MathWorks UK, Cambridge). MATLAB code of the models and Supplementary figures are presented in the Supplementary Information file.

### 3.2. Experimental methods

The assembly reactions was done as described before (10). Briefly, we used a 2 µM aliquot of ϕC31 integrase and final concentrations were 3 nM vector (*BB*/pSira1 or *PB/* pSira3) and 5 nM PCR fragments (unless stated otherwise). Percentage of correct colonies (red) was taken from streaks of individual colonies to make the colour clearer. Total reaction time was 6 h unless stated otherwise. All reactions were ethanol-precipitated and the precipitated DNA was used to transform electrocompetent cells. The relationship of the number of colonies to known concentrations of the final product is shown in Fig. 11.

**Figure 11.**
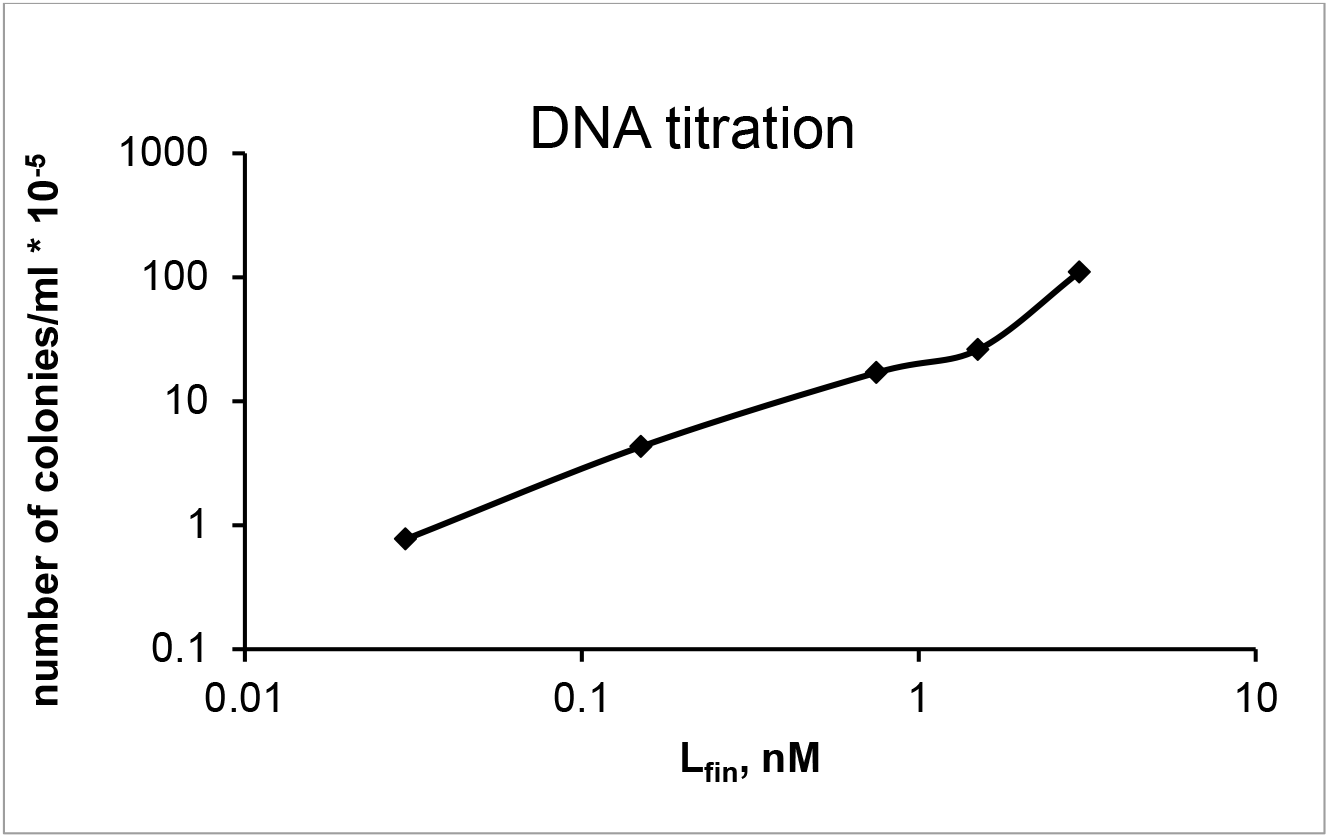
Titration of the number of colonies against known concentration of the final product of 3-fragment *PP/BB* assembly.

## 4. Conclusions

To analyse and improve methods for DNA fragment assembly with serine integrases, we built a mathematical model. The model explains previous observations and provides an instrument for optimization of assembly. The reduction of yield with increased number of fragments is explained by accumulation of unproductive synaptic complexes between mismatched sites. Increased amounts of wrong products (*wp*) with more fragments is explained by slower rate of *in vitro* inactivation of integrase, due to its protection by higher DNA levels. The combined effects of more mistakes and lower yields explains the observed sharp increase of the proportion of wrong products with more fragments. The model predicts that increased yields and reduction of mistakes can be achieved by increasing the fragment concentrations and decreasing the vector concentration. Assembly should avoid using substrates that stimulate the formation of *wp*. In particular, the use of a vector with mismatched *P* and *B* sites, and fragments leading to single-step formation of wrong products, should be avoided.

To test the model predictions and explore the mechanisms of *wp* formation, we performed experiments on 1- and 3-fragment assemblies. We confirmed that integrase inactivation during the course of the reactions has a noticeable effect on assembly kinetics. We also confirmed that assembly might be improved by increase of fragment concentrations and decrease of the vector concentration. Our data further suggest that mechanisms of ϕC31 integrase-mediated assembly might be more complex than is assumed by the model. Thus, reactions between mismatched sites might depend on the exact sequence of the central dinucleotides. Also, the use of assembly with mixed types of fragment (*PB, PP, BB*) might be complicated by intramolecular recombination between the mismatched sites of *PB* substrates and formation of long recombinant intermediates, depleting the substrates and leading to *wp*. More data are required to adjust the model parameters, which would allow us to use the model for further exploration. Additionally, assembly reactions might benefit from using more efficient integrases, such as Bxb1 integrase. The model can be easily re-fitted to other integrase data or extended for multi-integrase assemblies.

## Funding

Biotechnology and Biosciences Research Council grant BB/K003356/1.

5.

## 6. Appendix A

### 6.1 Equations of 1-fragment PP/BB assembly

The model equations for 1-fragment *PP/BB* assembly (Fig. 3A, Fig. S2A) are: 
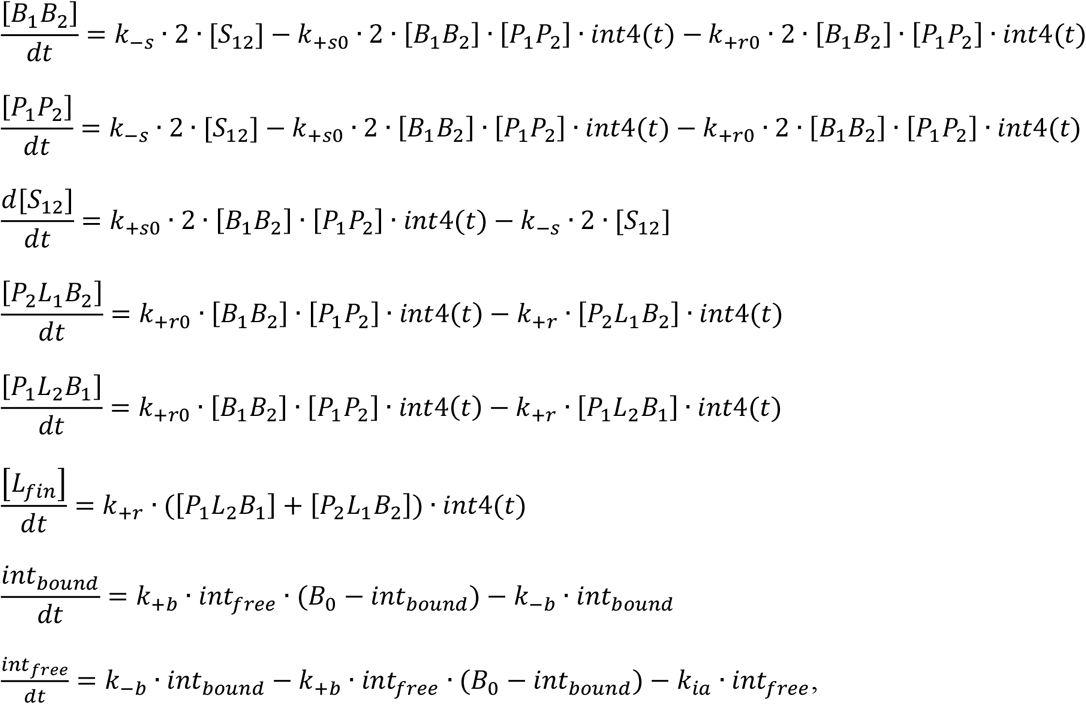
where *[B_1_B_2_], [P_1_P_2_], [P_1_L_2_B_1_], [P_2_L_1_B_2_]* and *[L_fin_]* are the concentration of free vector, fragment 2, *P_1_L_2_B_1_, P_2_L_1_B_2_* and the final *L_fin_* product, respectively (Fig. S2 A); *[S_12_]* is the concentration of *ws S_12_*; *int_free_* and *int_bound_* are concentrations of free and bound integrase; *B_0_* and (*B_0_ – int_bound_*) are the concentration of total and free sites for integrase binding; *int4(t)* is a function of integrase concentration: 
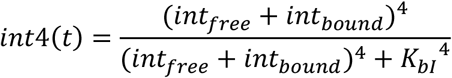

*B_0_ = k*·(*[B_1_B_2_]_0_*+*[P_1_P_2_]_0_*), where *[B_1_B_2_]_0_* and *[P_1_P_2_]_0_* are initial concentrations of the vector and fragment 2. All concentrations are expressed in µM; the time unit is min.

### 6.2. Equations of 1-fragment PB assembly

The model equations for 1-fragment *PB* assembly (Fig. 3B, Fig. S2B) are: 
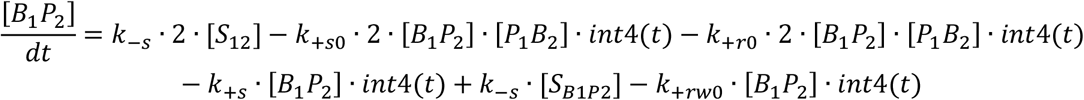

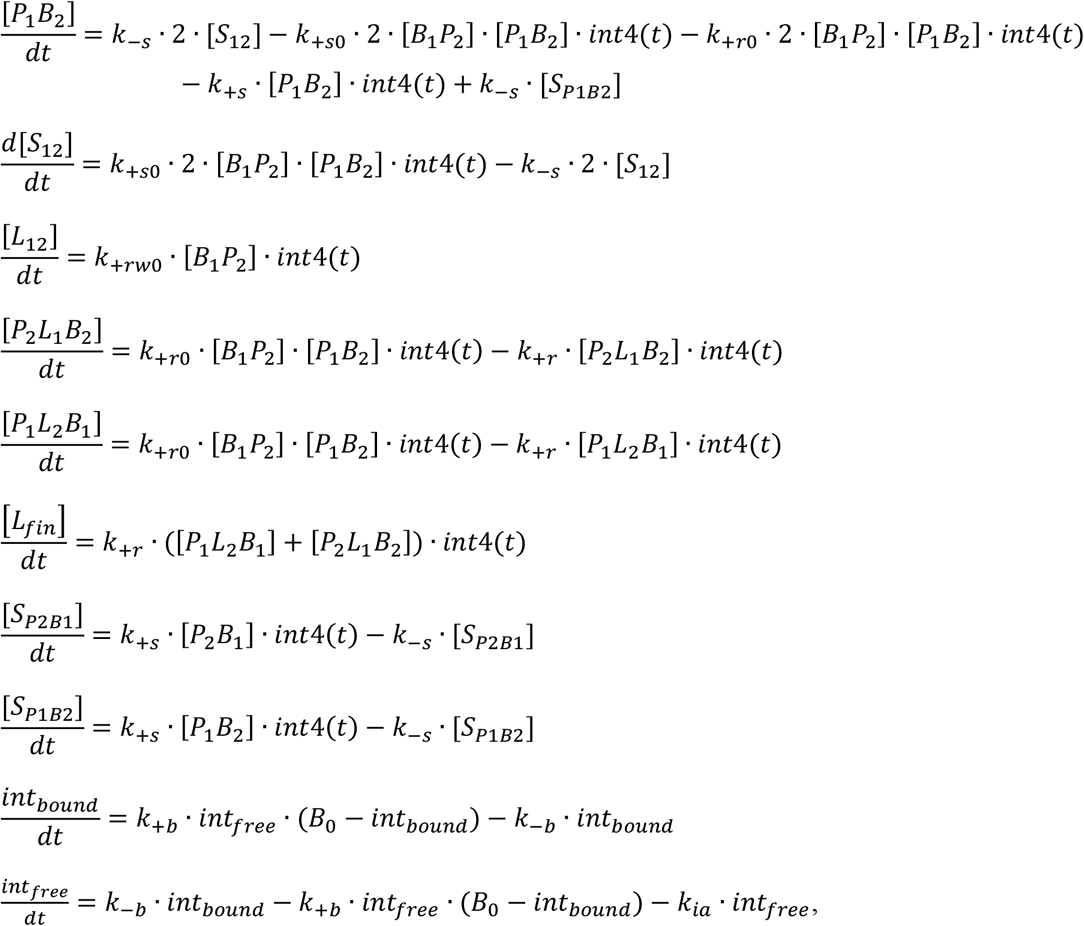
where *[B_1_P_2_], [P_1_B_2_], [P_1_L_2_B_1_], [P_2_L_1_B_2_]* and *[L_fin_]* are the concentration of free vector, fragment 2, *P_1_L_2_B_1_, P_2_L_1_B_2_* and the final *L_fin_* product, respectively; *[S_12_], [S_B1P2_], [S_P1B2_]* are the concentrations of *ws S_12_, S_B1P2_, S_B2P1_*; *[L_12_]* is the concentration of *wp L_12_* (Fig. S2 B); *int_free_* and *int_bound_* are concentrations of free and bound integrase; *B_0_* is the total concentration of integrase binding sites; *int4(t)* is a function of integrase concentration: 
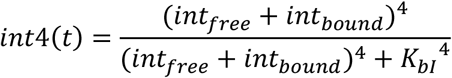
*B_0_ = k*·(*[B_1_P_2_]_0_*+*[P_1_B_2_]_0_*), where *[B_1_P_2_]_0_* and *[P_1_B_2_]_0_* are initial concentrations of the vector and fragment 2. All concentrations are expressed in µM; the time unit is min.

### 6.3. Equations of 3-fragment PP/BB assembly

The model equations for 3-fragment *PP/BB* assembly (Fig. 3C, Fig. S2 C) are: 
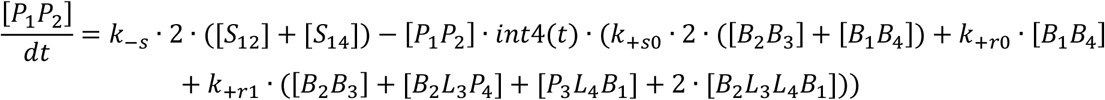

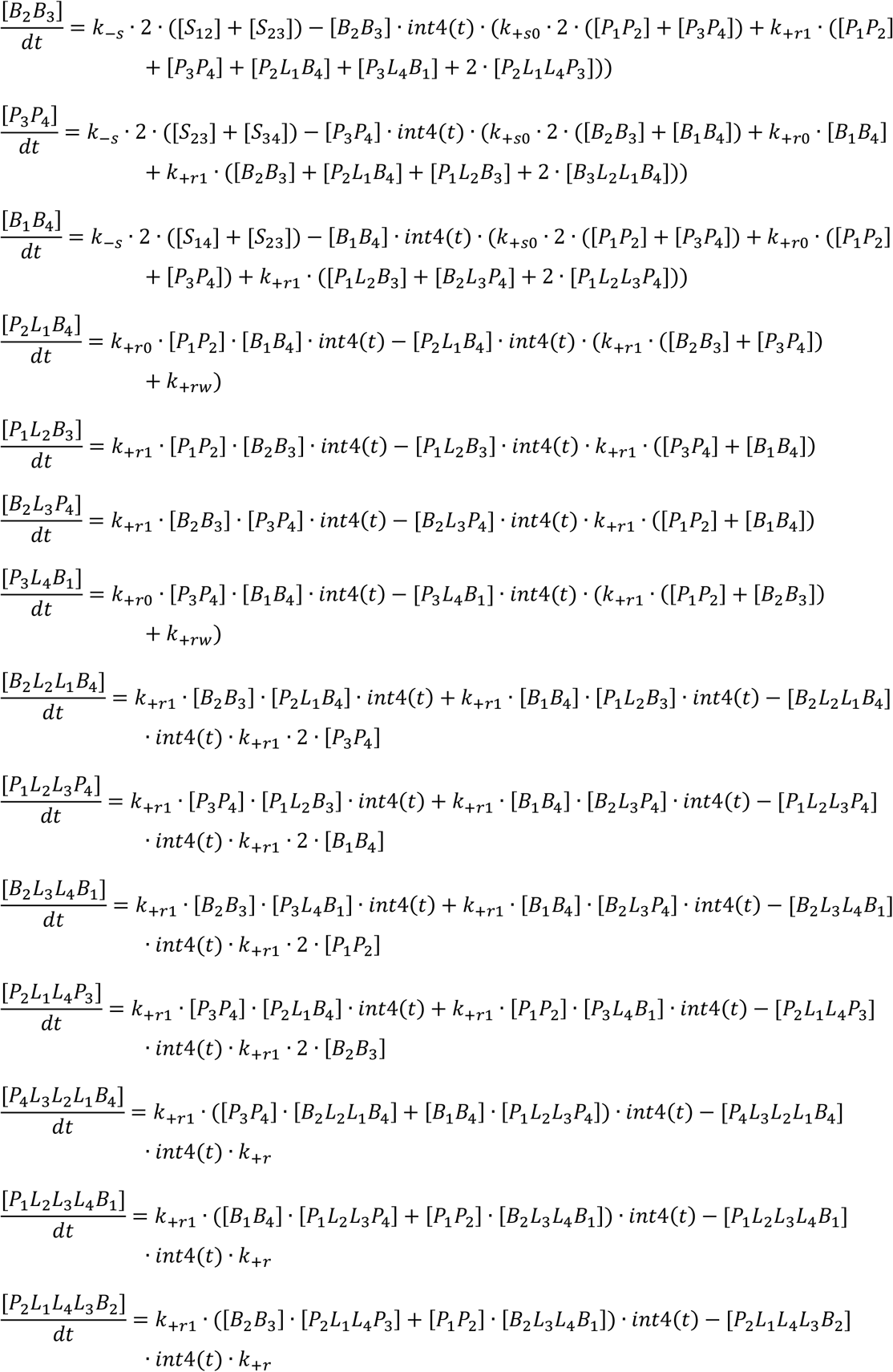

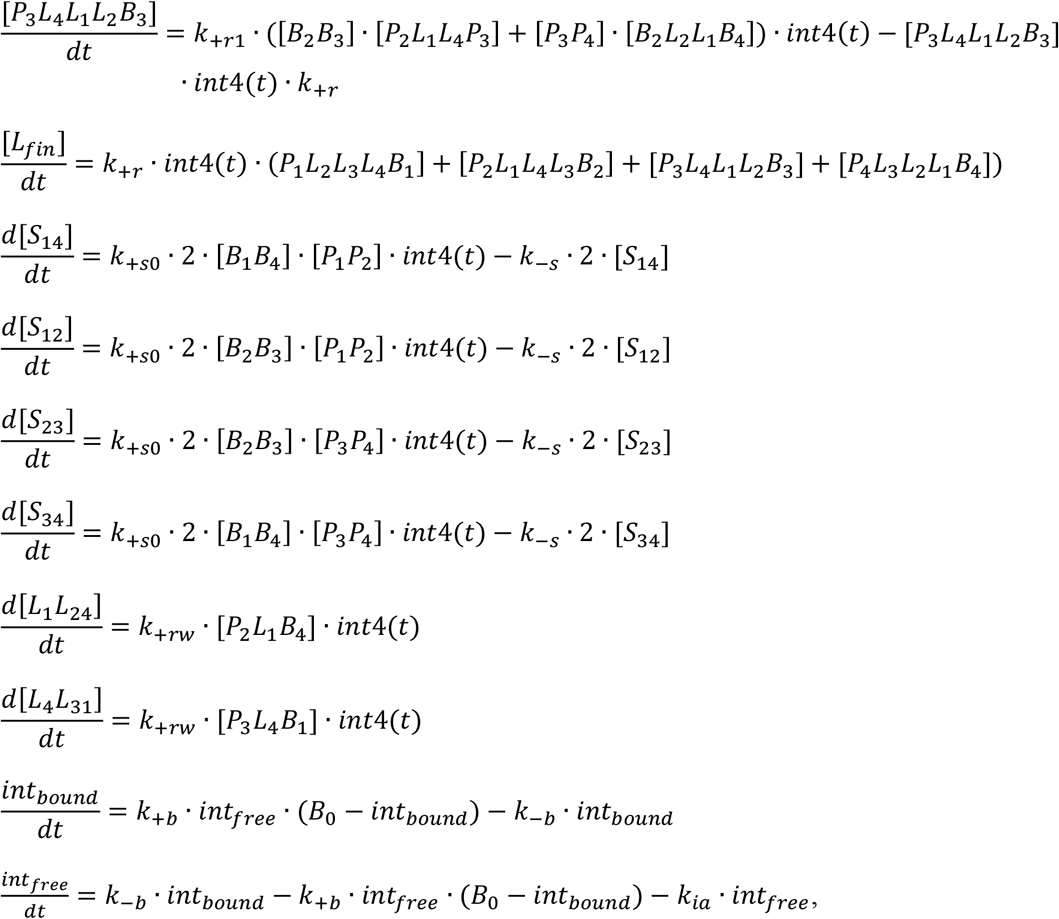

Where *[P_1_P_2_], [B_2_B_3_], [P_3_P_4_], [B_1_B_4_]* and *[L_fin_]* are the concentration of free fragments 1, 2, 3, vector and the final *L_fin_* product; *[P_2_L_1_B_4_], [P_1_L_2_B_3_], [B_2_L_3_P_4_], [P_3_L_4_B_1_], [B_3_L_2_L_1_B_4_], [P_1_L_2_L_3_P_4_], [B_2_L_3_L_4_B_1_], [P_2_L_1_L_4_P_3_], [P_4_L_3_L_2_L_1_B_4_], [P_1_L_2_L_3_L_4_B_1_], [P_2_L_1_L_4_L_3_B_2_], [P_3_L_4_L_1_L_2_B_3_]* are the concentrations of the respective recombinant intermediates; *[S_12_], [S_14_], [S_23_], [S_34_]* are the concentrations of *ws*: *S_12_, S_14_, S_23_, S_34_*; *[L_1_L_24_]* and *[L_4_L_31_]* are the concentrations of *wp*: *L_1_L_24_* and *L_4_L_31_* (Fig. S2 C); *int_free_* and *int_bound_* are concentrations of free and bound integrase; *B_0_* is the total concentration of integrase binding sites; *int4(t)* is a function of integrase concentration: 
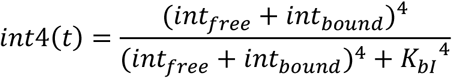

*B_0_ = k*·(*[P_1_P_2_]_0_*+*[B_2_B_3_]_0_*+*[P_3_P_4_]_0_*+*[B_1_B_4_]_0_*), where ·*[P_1_P_2_]_0_, [B_2_B_3_]_0_, [P_3_P_4_]_0_* and *[B_1_B_4_]_0_* are initial concentrations of fragments 1, 2, 3 and the vector. All concentrations are expressed in µM; the time unit is min.

## References

1. Brophy JA, Voigt CA. Principles of genetic circuit design. Nature Methods. 2014;11(5):508–20.

2. Shih PM, Liang Y, Loque D. Biotechnology and synthetic biology approaches for metabolic engineering of bioenergy crops. Plant J. 2016;87(1):103–17.

3. Johns NI, Blazejewski T, Gomes AL, Wang HH. Principles for designing synthetic microbial communities. Curr. Opin. Microbiol. 2016;31:146–53.

4. Olorunniji FJ, Rosser SJ, Stark WM. Site-specific recombinases: molecular machines for the Genetic Revolution. The Biochemical Journal. 2016;473(6):673–84.

5. Gibson DG. Synthesis of DNA fragments in yeast by one-step assembly of overlapping oligonucleotides. Nucleic Acids Res. 2009;37(20):6984–90.

6. Engler C, Kandzia R, Marillonnet S. A one pot, one step, precision cloning method with high throughput capability. PloS One. 2008;3(11):e3647.

7. Fogg PCM, Colloms S, Rosser S, Stark M, Smith MCM. New applications for phage integrases. J Mol. Biol. 2014;426(15):2703–16.

8. Baltz RH. *Streptomyces* temperate bacteriophage integration systems for stable genetic engineering of actinomycetes (and other organisms). Journal of Industrial Microbiology & Biotechnology. 2012;39(5):661–72.

9. Venken KJT, He Y, Hoskins RA, Bellen HJ. P[acman]: a BAC transgenic platform for targeted insertion of large DNA fragments in *D. melanogaster*. Science. 2006;314(5806):1747–51.

10. Colloms SD, Merrick CA, Olorunniji FJ, Stark WM, Smith MCM, Osbourn A, et al. Rapid metabolic pathway assembly and modification using serine integrase site-specific recombination. Nucleic Acids Res. 2014;42(4):e23.

11. Smith MCM. Phage-encoded serine integrases and other large serine recombinases. In: Craig NL, Chandler M, Gellert M, Lambowitz AM, Rice PA, Sandmeyer S, editors. Mobile DNA III. 3 ed. Washington, DC: ASM Press; 2015. p. 253–72.

12. Pokhilko A, Zhao J, Ebenhöh O, Smith MC, Stark WM, Colloms SD. The mechanism of ϕC31 integrase directionality: experimental analysis and computational modelling. Nucleic Acids Res. 2016.

13. Smith MCA, Till R, Brady K, Soultanas P, Thorpe H, Smith MCM. Synapsis and DNA cleavage in ϕC31 integrase-mediated site-specific recombination. Nucleic Acids Res. 2004;32(8):2607–17.

14. Olorunniji FJ, Buck DE, Colloms SD, McEwan AR, Smith MC, Stark WM, et al. Gated rotation mechanism of site-specific recombination by ϕC31 integrase. Proceedings of the National Academy of Sciences of the United States of America. 2012;109(48):19661–6.

15. Fan HF, Hsieh TS, Ma CH, Jayaram M. Single-molecule analysis of ϕC31 integrase-mediated site-specific recombination by tethered particle motion. Nucleic Acids Res. 2016;44(22):10804–23.

16. Ghosh P, Kim AI, Hatfull GF. The orientation of mycobacteriophage Bxb1 integration is solely dependent on the central dinucleotide of attP and attB. Mol. Cell. 2003;12(5):1101–11.

17. Shore D, Langowski J, Baldwin RL. DNA flexibility studied by covalent closure of short fragments into circles. Proceedings of the National Academy of Sciences of the United States of America. 1981;78(8):4833–7.

